# Linguistic contextualization in the human hippocampus

**DOI:** 10.1101/2025.06.23.661103

**Authors:** Kalman A. Katlowitz, James L. Belanger, Taha Ismail, Ana G. Chavez, Assia Chericoni, Melissa Franch, Elizabeth A. Mickiewicz, Raissa K. Mathura, Danika L. Paulo, Eleonora Bartoli, Steven T. Piantadosi, Nicole R. Provenza, Andrew J. Watrous, Alica M. Goldman, Vaishnav Krishnan, Atul Maheshwari, Sameer A. Sheth, Benjamin Y. Hayden

## Abstract

Word meanings in language are contextualized by surrounding words. Inspired by the self-attention mechanism in transformer-based large language models (LLMs), we hypothesized that structural composition in the brain arises from combining canonical (non-contextual) word representations with those of nearby words. We analyzed single unit activity in the human hippocampus, a region involved in semantic and contextual processing, while n=10 participants listened to podcasts. We found that hippocampal neurons encoded word position within a clause, using both ordinal and frequency-domain positional encoding. Moreover, neural responses to specific words reflected both the word’s own lexical semantics and a weighted sum of the embeddings of preceding words. The relative weighting of these contextualizing words correlated with LLM self-attention weights. These findings suggest that contextualization in the brain makes use of vectorial shifts that have a resemblance to attentional reweighting in LLMs, and highlight the role of the mesial temporal lobe within the broader language network.

## INTRODUCTION

If words are the building blocks of language, contextualization is its architecture^1–3^. Contextualization includes, for example, adverbial modification (“*very tall*”), long-distance dependencies like pronoun reference (“*the dog chased its tail*”), adjectival composition (“*red balloon*”), and prepositional marking (“*up the mountain*”). In linguistic theories, contextualization in LLMs has perhaps its closest analogue in dependencies, where words have specific relationships with other words^4,5^. Generally speaking, contextualization encompasses multiple types of dependency, as indicated by the above examples. Impairments in the ability to understand contextualization are hallmarks of schizophrenia, autism, and other psychiatric diseases^6,7^, and contextualization is an important element of not just speech, but many features of higher-level human cognition^8,9^.

Large Language Models (LLMs) leverage large training sets and machine learning principles to replicate key features of human language use^10–14^. Modern LLMs start with a dictionary, wherein each word (or sub-word token) is represented by a vector of numbers. They then implement contextualization through *self-attention*, by which they use learned associations between specific word pairs to combine basic word representations together into new vectors that represent their semantic meaning within a specific context (this process includes both linear and non-linear processes, see **Discussion**^15,16^). The magnitudes of these weighted combinations vary on a word-by-word basis, so that contextualization is graded, rather than all-or-none. To implement contextual weightings, LLMs make use of positional encoding, which represents the locations of words within the larger clause^17^. Ultimately, LLMs’ weights are optimized for next-word prediction, which is sufficient to drive the learning of contextual relationships and results in internal representations of key structures and dependencies of language. Self-attention, therefore, instantiates a specific theory of contextualization, one with testable hypotheses for how the human brain may implement it.

Classic models of the neural basis of contextualization focus on *where* rather than *how*. That is, they generally focus on questions of functional neuroanatomy^18–20^, putting aside neurocomputational mechanism (though see refs 2,21–24). It is well documented, however, that neural representations of words are highly modified by the local context. For example, semantic contextualization improves decodability^25–27^. Indeed, there is evidence for reactivation of specific adjective semantic responses during listening to subsequent nouns they modify^28–30^. Inspired by transformer architecture models in LLMs, we hypothesized that contextualization makes use of weighted summation of semantic embedding vectors^31^, which in turn correspond to semantic encoding across neurons in the human brain^32–36^.

We are especially interested in the hippocampus, a region that is responsible for encoding general semantic information^37,38^, and whose single units show distributed encoding of semantics^32,39,40^. The hippocampus tracks higher order features of speech, such as grammatical categories^28,41,42^. Although it is not part of the canonical language network^8^, the hippocampus consistently engages in language comprehension, especially word meanings^43–47^ and plays a critical role in contextualization across multiple functional domains^48^. Indeed, patients with hippocampal damage often exhibit deficits in distinguishing similar experiences that occur in different contexts^49–53^ including linguistic ambiguities^54^.

To test the hypothesis that the hippocampus is engaging in semantic contextualization, we examined hippocampal single neuron responses during passive podcast listening. We found broad support for the idea that the hippocampus recapitulates three key elements of self-attention transformer processes. ***First***, neurons in the human hippocampus show clear positional encoding, including both a multi-dimensional linear scheme and a sinusoidal (Fourier) scheme. ***Second***, contextual responses to words reflect a vector sum of the features defining the word, i.e. neural responses to words are influenced by weighted combinations of prior contextualizing words. ***Third***, the weighting of these contextual words reflects the weightings associated with those words used in LLMs. ***Finally***, we demonstrate that neuronal firing patterns reflect not just the semantic information of the current word in isolation, but also a combination of prior words that contextualize it. Together these results support the idea that the brain and LLMs share a common set of computational principles in the implementation of semantic contextualization and advance our understanding of the mechanisms underlying speech comprehension.

## RESULTS

We recorded responses of 356 isolated stable units in the hippocampus of ten native English-speaking patients who were undergoing neural monitoring for epilepsy. Each patient listened to the same 47 minutes of English speech from a popular podcast (six monologues taken from the Moth Radio Hour), comprising a total of 7346 words for each patient (**Figure 1A**; **Methods**). An analysis of the dataset used here was previously reported in a study demonstrating semantic encoding in the hippocampus^32^. For example, individual units can differentiate between nouns and verbs (55.9% of units, rank sum test, alpha =0.05), social vs. nonsocial words (46.6%, rank sum test, alpha =0.05), and concrete versus abstract words (3.7%, Spearman’s correlation, applying per neuron familywise (Benjamini-Hochberg) error rate correction for all). Here, we examined how neuronal responses to words change as a function of context (**Figure 1B**).

**Figure 1.**
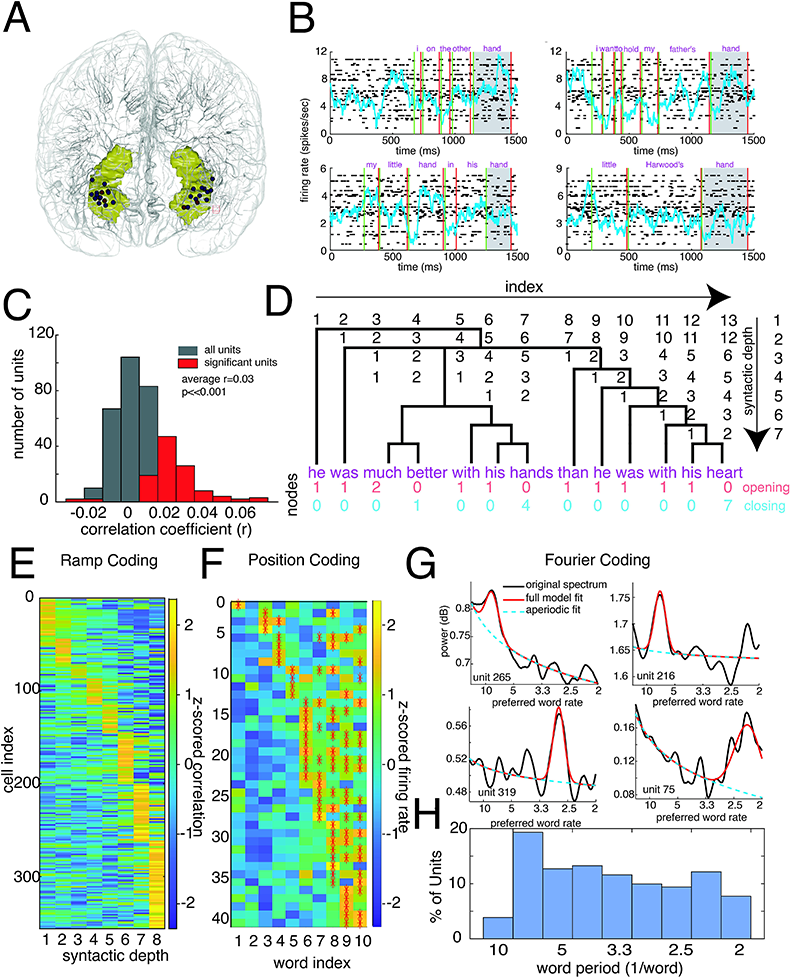
Human hippocampal neurons encode relative word positions during a passive listening task. **A.** Electrode recording location of hippocampal neurons from ten patients, red sites refer to the data obtained in B. All brain renders created with code from ref. 109. **B.** Population response of 54 units in a single patient to four different phrases, each of which had a different meaning of the word “hand.” Green lines are onsets, and red lines are offsets of each word. **C.** Distribution of correlations between firing rate and word index in a sentence for all (blue) and significant (red) units **D.** Diagram of hierarchical indexing as a function of syntactic depth. The number of nodes opening and closing are listed below in red and blue, respectively. E. Z-scored correlation values across each syntactic depth for each unit sorted by the index of the highest z score. F. Z scored mean firing rate of units significant for any index as a function of index within a sentence for the first ten positions. Red asterisks indicate significance on rank sum test. G. Example power spectra (black) of the word level firing rates of significant units, with the dominant peak (red) highlighted above the aperiodic component (dotted blue line). H. Distribution of center frequencies for all significant units demonstrating tiling across the frequency space.

### Positional encoding by hippocampal neurons

No matter the mechanism for semantic contextualization, it can be helpful for the brain to track the ordinal relationships between words. For example, despite having an identical set of words, the phrases “*the dog chased the man*” and “*the man chased the dog*” have different meanings. We first examined how word position could be represented within the hippocampus. For each neuron we defined the word response as the average firing rate during the playing of that specific word (adding an 80 ms lag), and assessed the role of word position on that response.

#### Ramp Coding

We found a significant correlation between a neuron’s response to each word and that word’s position within a sentence in 31.7% of units (n=113/356, α<0.05, Kendall’s rank correlation). The majority of significant correlations were positive (96.4% of the 109 significant units; 29.5% of all units) and correlations were low overall, average r=0.03 +/-0.01 (Spearman’s correlation, mean +/-standard deviation, **Figure 1C**). These results were reproducible from a decoding approach as well (r^2^ = 0.015 on held out data, ridge regression, 10 fold cross validation, p<0.01), suggesting that approximately 1.5% of the variance can be explained by ramp coding. This overall bias towards an increase in firing rate for words later in a sentence is reminiscent of previous studies showing an increase in spectral power across word index, which was attributed to working memory load^55,56^. Although we cannot fully disambiguate the effects of position and memory load, we can use correlates of phrasal - and thus language-specific - working memory complexity, such as opening nodes or closing nodes^57,58^ (see **Methods**). Opening nodes correspond to linguistic states that enable continued information flow (“he was <u>much</u>…”), whereas closing nodes mark points at which information flow is terminated (“with his <u>heart</u>” **Figure 1D, bottom**). We created a linear model regressing firing rate against all three parameters simultaneously; we still found a strong residual correlation between index and firing rate in 18.0% of units (n=64/356), versus number of closing nodes (6.5%, 23/356) or opening nodes (12.9%, 46/356), consistent with a main effect of position. This result was also maintained when controlling for other covariates of word index, such as mean dependency distance, part of speech and word frequency (15.5% of units significant, with 83.6% of those 55 units having a positive relationship to word index).

Sentences are characterized by structure at multiple levels, and a word may have different positions at different levels (e.g., the first word in a dependent clause can be in the middle of the sentence). We examined positional encoding separately at each of eight increasing syntactic depths (depths calculated using CoreNLP^59^, see **Methods**; **Figure 1D**). We found neurons with preferential encoding of position at each depth (**Figure 1E**). Firing was more likely to be modulated by the indices of deeper syntactic levels (χ(5)^2^ =76.7, p<0.0001, correlation between depth and number of significant units, r =0.74, p<0.05). To account for correlations in word index across syntactic depths, we ran a multivariate regression to predict firing rate as a function of indices at all depths. Consistent with the preference towards deeper syntactic levels found in the univariate analysis, the median absolute value of the coefficients increased as a function of syntactic depth (r=0.89, Spearman’s correlation, p=0.003, **Figure 1F**).

We next examined other features that could modulate positional effects. To quantify the impact of sentence-level semantics, we used the first 10 PCA dimensions of the final pooled embedding of the General Text Embeddings (GTE) sentence level language model^60^. In a model that incorporated position, GTE parameters, and their interactions we still found a significant number of units that were significant purely for position (17.7% of units, t-test on the coefficient). However, there were also units that were significant for interactions with position (12.4%, F-test at least one significant interaction term), often the same units (χ(1)^2^ =4.8, p=0.03). We also examined the impact of sentence length and syntactic complexity. Here too, sentence position was still the strongest regressor, both in terms of number of significant neurons (16.8%) and effect size (0.11+/-0.03, mean +/-SEM). Interaction effects were also significant at 10.6% and 11.2% of units at negative (−0.06+/-0.01) or mixed (0.007 +/-0.008) directions for syntactic depth and sentence length, respectively (**Extended Data Fig. 1 A-D**). Thus, while many different thematic and syntactic features are correlated with unit firing rate, sentence position still has the most salient impact.

To dissociate between a syntactic response and a semantically specific tracking of position we then analyzed a separate dataset of Jabberwocky stimuli (7 patients, 272 units, 1797 words), which alters words into nonsense while preserving syntactic structure^61^ (see **Methods**). While some neurons still tracked position, the incidence (12% of units) was smaller than for language data (p<0.0001, Fisher’s exact test), as was the mean significant effect size (r=0.02 +/-0.007, Wilcoxon rank sum test z=2.8, p=0.005) (**Extended Data Fig. 1EF**). A subset of units were recorded across both tasks (3 patients, 85 units). While these units’ propensity to encode position was similar in both tasks (r=0.23 p=0.03, Spearman’s correlation), the encoding was stronger in the podcast task (**Extended Data Fig. 1G**, Wilcoxon signed rank test z=2.7, p=0.007). As such, while neurons can have increased firing with position in sentence even without any normal sentence semantics, positional encoding is strengthened when semantic meaning is involved.

#### Index Coding

A different (but not mutually exclusive) encoding scheme is selectivity for a specific word index. For example, in the premotor nucleus of the zebra finch, a population of narrowly tuned projection neurons are each selective for a specific time point and together represent relative position within a song sequence^62^ (see also ref. 63 for other animal models). In other words, instead of an overall change in firing across word numbers, neurons respond preferentially to a specific word number, such as the third word in a clause. We found index selectivity for a modestly sized population of units across multiple syntactic depths (13.2 +/-4.5% of units, Kruskal-Wallis test on indices 1-10, α<0.05). However, unlike in the zebra finch HVC, these neurons were typically selective for more than one word index (e.g. sentence level, **Figure 1G**); they had multiple positions significant on the post-hoc rank sum test for each index versus all others (1.8 +/-0.2 significant positions per cell, proportion with one peak: 37.7 +/-9.5%, chance at 38.7%). The existence of multiple peaks raises the possibility of sinusoidal coding.

#### Sinusoidal coding

A third (also non-exclusive) positional encoding scheme is used by some LLMs, including in Rotary Position Embedding (RoPe^16,64^). Sinusoidal positional encoding has the advantage of scalability to long clauses. Sinusoidal positional encoding would show up as a consistently modified response to words occurring at specific periods, such as an increase in firing rate for every other word or every fourth word. To test for this code, we computed the power spectrum of the firing rate response to each word across the recording and detected peaks above the aperiodic component^65^ for each neuron (**Figure 1H**). Despite a lack of any intrinsic periodicity within our task, we found many instances of oscillatory tuning, with 50.8% of units (n=181/356) showing a preferred word phase (α=0.05, peak greater than shuffled index power spectrum peak). Said differently, about half the neurons showed modulation in their firing rates once every *n* words, where n is ranging from 2 (every other word) to 20. These units were not identical to the ramp (χ(1)^2^=0.11, p=0.7) or index (χ(1)^2^=1.4, p=0.2) units. Moreover, their anatomical position was slightly more medial, anterior, and inferior (p<0.05 for all, rank sum test) than the ramp or index coding units.

Across the population, peak frequencies approximately tiled the frequency space (**Figure 1H**, Kolmogorov–Smirnov (KS) test versus uniform distribution D=0.08, p=0.2), as would be predicted by an efficient code. This frequency tuning was not stable over the course of the experiment (e.g. **Extended Data Fig. 2A**, p=0.2, Spearman’s correlation between first and second half), but there was preservation of both the identity of the frequency representative units (χ(1)^2^ =27.0, p<0.001 **Extended Data Fig. 2B**) and the uniformity of the frequency representation (**Extended Data Fig. 2C,D**, p>0.05 KS test for all except episode 4, D=0.12, p=0.002 overrepresenting longer word periods). As such, there is a stable population of neurons involved in signalling word order in the frequency domain, although their tuning remaps across contexts.

This apparent sinusoidal encoding could be an artifact of neurons having an intrinsic (non-stimulus driven) firing rate oscillation at a fixed temporal frequency. To rule out this alternative explanation, we tested for any intrinsic periodicity for the neurons during the rest periods before and after the task (62.0 +/-10.1 total minutes per patient) and compared the intrinsic periodicity to preferred word periods (range of 0.45 to 2.25 Hz, corresponding to a median word rate of 4.5 words per second). Of the units with a preferred word period, 34% (61/181) had no significant undriven oscillatory power (16.7% of total population). However, even among the inherently oscillatory neurons, we observed no systematic relationship between their preferred undriven period and the preferred word period (r=0.04, p=0.7, Spearman’s correlation), indicating that lexical sinusoidal positional encoding is not simply a by-product of intrinsic periodicity.

There are multiple lines of evidence suggesting there is a functional difference between anterior and posterior hippocampus^66^. However, we found no difference between anterior and posterior neurons (**Extended Data Fig. 3A**) in the degree of ramp coding (p=0.8, rank sum test on the number of significant syntactic depths per unit, **Extended Data Fig. 3B**), proportion of neurons that showed selectivity to index encoding (**Extended Data Fig. 3C**), nor the proportion of units that had a significant peak in the Fourier domain (**Extended Data Fig. 3D**, Fisher test p=0.4) or their center frequency **(Extended Data Fig. 3E**, two sample t-test, t(243)=0.98, p=0.3).

### Hippocampal neurons encode attentionally weighted sums of semantic vectors

We next asked how neuronal responses are modified by the semantics of adjacent words. Inspired by self-attention models in LLMs, we hypothesized that neuronal responses might represent a weighted sum of semantic embedding vectors^16^ (**Figure 2A**). We defined canonical responses to each unique word as the average vector across all instances of that word. Next, the shifted response for individual words was then simply the deviation from that average. We can then measure the influence of prior words by assessing the alignment of their semantic vectors with this shift. To quantify this for each word, the relative pairing score of prior words was defined as Z, the z-score of these values across the antecedent 20 words. Consider for example, a sentence from one of our podcasts: “*I should probably tell you that the people he’s talking about are the football team.*” In this sentence, the neural response (a 356-dimensional vector corresponding to 356 neurons across 10 patients) evoked by the word “*team*” is shifted from its standard response (estimated from the average response evoked by the other four times it was presented). The contextualized response to “*team*” is directed towards the word “*football*”, meaning that it can be represented as a weighted sum of the non-contextual neural responses for “*team*” and “*football*” (**Figure 2B**). By this measure, the second most strongly contextualizing word was “*people*”. Meanwhile, the word “*he’s*” has the lowest similarity, presumably because the character it refers to is being contrasted with the football team. Averaged across all words in our stimulus set, we find that words are contextualized as a function of relative distance, i.e. they are more sensitive to the words that immediately precede them (**Figure 2C**).

**Figure 2.**
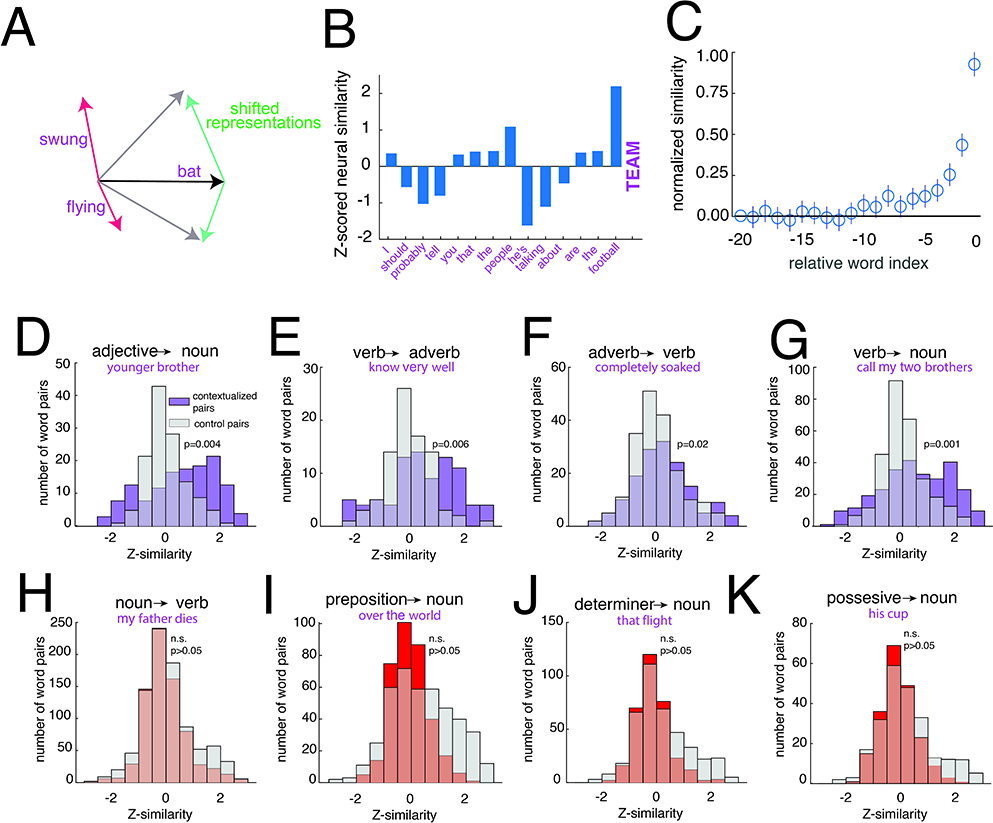
Semantic relationships are represented through contextually weighted vector summations. **A.** Vectorial diagram of how adjacent words can modify the neural vectorial representations of words (e.g. ”swung” appearing in the context of “bat” shifts the firing rate vector for “bat” to have a more similar cosine distance to the vector for “swung”) **B.** Z-scored cosine similarity values between the shifted neural representation of the final word “team” and all preceding words in the sentence. **C.** Normalized (0 to 1) average z-scored cosine similarity values as a function of relative index. n=7346 words. Error bars represent +/-2*SEM, **D-K.** Distribution of the z-scored contextual pairing values for the word pairs designated above (purple/red) and the values for control word pairs (gray). Purple distributions were determined to be significantly different per signed rank test (see results).

To quantify specific word-type effects across our dataset, we investigated several basic grammatical pairings^59^. Note that for this analysis, words could have a significant pairing strength simply due to temporal autocorrelations or local context. Therefore, rather than using random word pairs as controls, we defined controls as pairs with the exact same interword distance by shifting all selected indices by one word earlier. We found that responses to nouns are consistently rotated in the direction of their antecedent adjectives (e.g. “**younger** *brother*”, n=137 adjective-noun pairs, Z=0.65+/**-** 0.13, right-tailed paired signed rank test versus nearby word pairs with a similar distribution of pairwise distances z=2.7, p=0.004, **Figure 2D**). Note that in this analysis, the antecedent adjective does not necessarily immediately precede the word it modifies (e.g. “**coolest** and meanest *kids*”). Our task did not include a sufficient number of samples to test post-nominal adjectives (n=4). Verbs contextualized adverbs that followed them (e.g. “**know** very *well*” n=86 verb-adverb pairs, Z=0.52+/**-**0.15, signed rank z=2.5, p=0.006 **Figure 2E**), just as adverbs contextualized the verbs that followed them (e.g. “**completely** *soaked*” n=158 adverb-verb pairs, Z=0.24 +/**-**0.09, signed rank z=2.0, p=0.02, **Figure 2F**). We also found that verbs contextualize the responses of nouns that follow them (typically, direct objects of the verb, e.g., “**call** my two *brothers*”, n=289 verb-noun pairs, Z=0.56+/**-**0.09, signed rank z=3.0, p=0.001, **Figure 2G**), but nouns (typically subjects) do not measurably contextualize the responses of verbs that follow them (e.g., “my **father** *dies*”, n=747 noun-verb pairs, Z=0.02 +**/-**0.03, signed rank z=-3.4 p>0.5, **Figure 2H**). Note that this pattern is consistent with the idea that heads contextualize tails more strongly than the reverse^26^ (see **Discussion**). Indeed, not all pairing types demonstrated a significant contextualization, perhaps due to a weaker semantic relationship or an abstract relationship. In prepositional phrases (“**over** the *world*”), the prepositions did not contextualize following nouns, despite a relatively large sample size (n=346, Z=**-**0.03 +/**-**0.03, signed rank z=-5.5, p>0.5, **Figure 2I**). Determiners also did not generally modify their respective nouns (“**that** *flight*”, n=324, Z=-0.13 +/**-**0.03, signed rank z=-4.4, p>0.5, **Figure 2J**); nor did possessive adjectives (“**his** *cup*”) contextualize their nouns (n=206, Z=**-**0.13 +/**-**0.04, signed rank z=-3.4, p>0.5, **Figure 2K**). The significant results were robust to the distance metric used, though with adverb:verb falling below significance when using Euclidean distance (**Extended Data Fig. 4A**). Note that for this analysis, the sign of the effect is meaningful. Thus, for example, adjectives typically have an overall positive impact on their nouns (**Extended Data Fig. 4C**, mean similarity 0.07 +/-.02, t-test t(136)=3.8, p<0.001) but words that are negated (e.g. **“not know”)** show a negative shift away from their average representations that approaches significance (**Extended Data Fig. 4D**, median similarity -0.1 +/-.04, signtest p=0.06).

### Correspondence between LLM contextual embedding and neural responses

The specific strength of mappings between the past and current words in transformer-based LLMs is given by the ***attention pattern grid*** (**APG**^67^) and is used by the LLM to define relationships. Note that while most LLMs have multiple attention heads whose effects summate, here we ignore this feature and focus instead on the total overall effect of attention. A high weight for a pair of words causes the embedding of the prior word to influence the representation of the current word. We reasoned that the projection of neural responses onto adjacent words (**Figure 2A, B**) allows us to infer what is in essence a **neural APG**. We next compared neural APGs to corresponding LLM-derived APGs. To generate the LLM-based APG, we treated GPT-2 dimension-wise responses as if they were unit responses and replicated the same process we used above to create an empirically derived estimator of the single APG for the LLM. For example, in our example sentence above, we find a close quantitative match between the neural data and the inferred APG from Layer 36 of GPT-2 (r=0.78 p<0.001, Spearman’s correlation, **Figure 3A**). Further, we find that the distributions of the word pairs’ contextual strength in an LLM **(Extended Data Fig. 5**) are similar to the neural data (**Figure 2D-K**).

**Figure 3.**
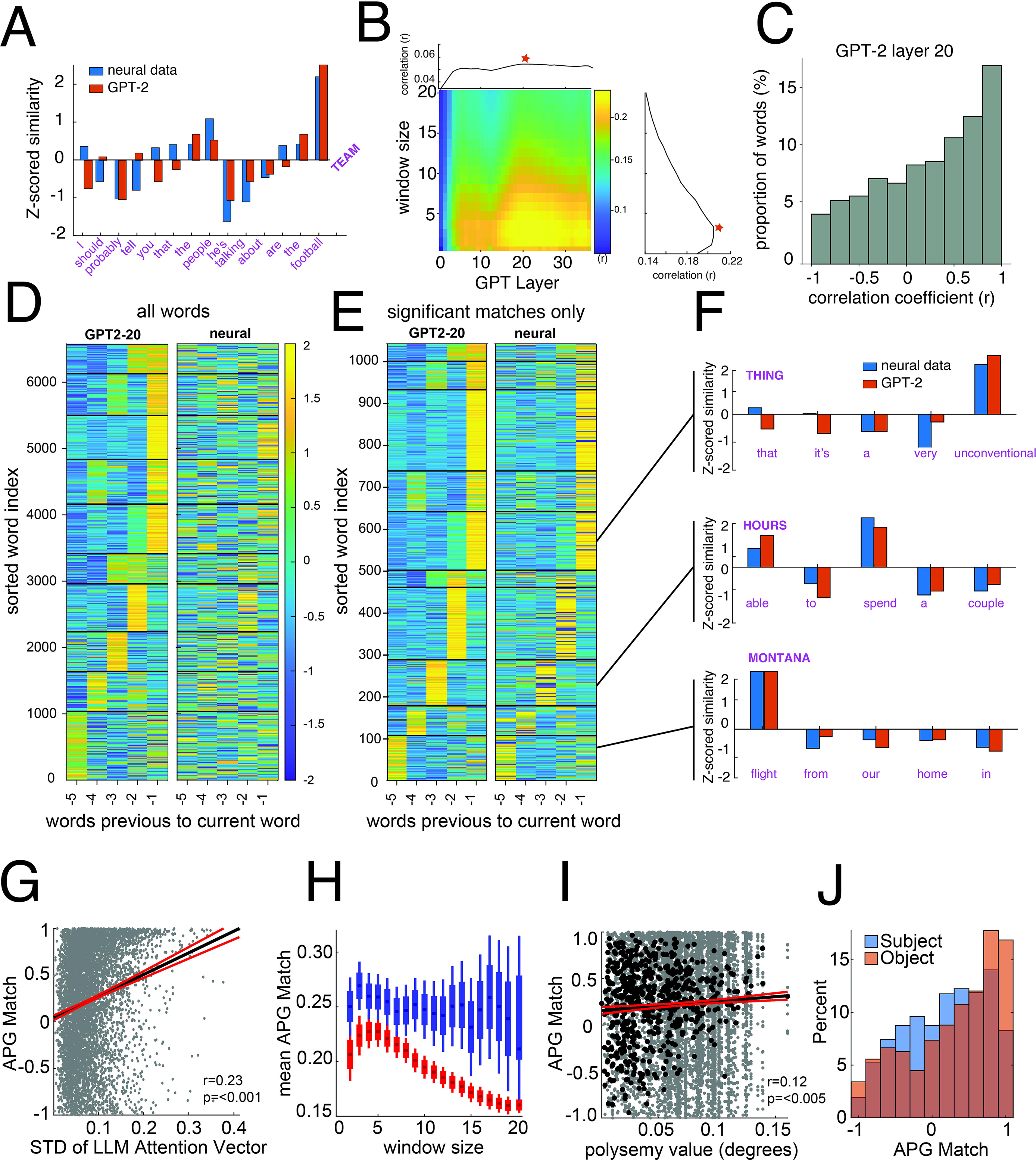
Neural attention patterns closely match LLM attention patterns. **A.** Similar to Figure 2B but with the APG vector for GPT-2 Layer 20 overlaid. **B.** Average match across all words between neural APG and GPT-2 APG for all layers (x axis) and window lengths from 2 to 20 (y axis). Average values projected above and to the right. Red star reflects the maximum. **C.** Distribution of r-values for correlations between the neural APG and the LLM embedding (GPT-2 Layer 20) APG for the previous 5 words for all words. **D.** The z-scored APG vectors for all words from GPT-2 layer 20 sorted by cluster index and peak (left) with the neural data sorted on the same indices (right). **E.** same as **C** but only words that had significant matches. **F.** Example phrases from the clusters. **G.** Spearman’s correlation between neural and LLM attention vectors as a function of the magnitude of the variance within the attention vector. Black and red lines represent the best fit +\- 95% confidence interval. **H.** Mean and SEM (box) and 2*SEM (whisker) of r values between the APG vectors of the LLM and the neural data for each word for all words (red, n range 6507 to 6581 words) and only words whose antecedent window is still within the same clause (blue, n range 104 to 5330 words). **I.** Spearman’s correlation between the match of the neural and LLM vectors of each word and the polysemy score of that word for all words (grey) and all unique words (black, r values are average across all presentations). Black and red lines represent the best fit +\- 95% confidence interval. **J.** Box and whisker plot representing the distribution of correlations between LLM and neural APG vectors for words that are subjects, objects, and prepositional objects.

We next created 37 latent LLM APGs, one for the initial embedding and then each of the 36 layers of GPT-2 and compared each one to our neural APG. To restrict our analysis to words likely to interact with each other (per **Figure 2C**) we tested the correlations across a range of local contextual windows (2-20) for each layer. We find that the mean correlation between neural and LLM APGs across words is positive for all layers and contextual windows, but can vary greatly as a function of both (**Figure 3B**). The correlation is maximized at Layer 20 and with a span of the five previous words (r=0.23 +/- 0.56, Spearman’s correlation p<0.001, **Figure 3C**). Note, however, that the overall fits were similar for sufficiently high layers, meaning that there is likely nothing special about Layer 20 compared to, for example, Layers 19/21 or even 15/25. Again our results were robust to the distance metric used, though peaking slightly earlier at layer 17 when using Euclidean distance (**Extended Data Fig. 4B**). Moreover, this result is not unique to GPT-2, which is decoder-based; we obtain similar results using the encoder-based LLM DeBERTa V3^68^ (maximized at layer 8 out of 12, r= 0.16 +/- 0.54, Spearman’s correlation p<0.001) and a different state-of-the-art decoder based LLM Llama-3^69^ (maximized at layer 18 out of 33, r= 0.24 +/-.07, Spearman’s correlation). We also find broadly consistent results across individual patients, with a bimodal peak in preferred layers and a preference towards smaller window sizes (**Extended Data Fig. 6**).

### A close quantitative match between neural and LLM-based APGs

We next examined the word-by-word temporal structures of the APGs (focusing on the Layer 20 and the five most recent words, see above) and identified ten distinct attention patterns using the Leiden algorithm (n=6579 words that all patients heard at least twice; **Figure 3D, left)**, a method that partitions the APGs into groups of words with similar temporal attention structure. While the immediately adjacent word was the most common strongest contextualizing influence, the LLM-based APGs demonstrated a striking variety of attention patterns. Remarkably, we find the same pattern of activity in the neural data when sorting by the same clusters, even though the LLM and neural data are completely independent of each other (**Figure 3D, right**). For example, a word with a high similarity index three words prior to it in LLM space tends to also have a higher similarity index three words prior to it in neural space. This effect is even clearer when only including words with a significant correlation between the neural and LLM attention vectors (n=1042 words, 16% of all words α=0.05, **Figure 3E**). A few examples demonstrating this phenomenon appear in **Figure 3F**, such as the phrase “*flight from our home in Montana*” representing the last cluster. Here, the prepositional object “*Montana*” is maximally modified by the subject “*flight*” occurring five words before it, and all other words in this cluster are maximally modified by the word five word prior.

What features of specific words predict that it will have a stronger match with the LLM-derived embedding? First, there was a strong relationship between the standard deviation of each word’s LLM attention vector and the subsequent match to the neural attention vector (r=0.21, p<0.0001, Spearman’s correlation, **Figure 3G**), suggesting that if a word’s meaning is not modified by prior words then the match will be poor simply because there are no features for our metric to capture^70^. Interestingly, despite finding a decay in the match with window sizes larger than the 5 prior words (**Figure 3B**, right), this effect was not found for word pairs that are still within a clause (**Figure 3H**), demonstrating that the hippocampus can dynamically adjust its “pairing memory” as a function of the dynamic grouping of words. Of note, the mean clause size was 4.7 +/-4.9 words, which likely accounts for the fact that our optimal window length is 5 words (**Figure 3C**).

Next, we examined the semantic features of the words that showed contextualized effects. For each word, we created a polysemy index, defined as the variance in its contextual embeddings across occurrences^32^. We found a positive correlation between this index and neural contextualization effect (r=0.05, p<0.0001, Spearman’s correlation), demonstrating that the more variable a word’s embedding, the better the neural definitions of relationships correspond to the LLM defined relationships. This effect size was stronger when averaging across all representations of each unique word (r=0.12, p<0.005, Spearman’s correlation, **Figure 3I**). This difference could also be seen within different grammatical subcategories: direct objects (that are defined by the verbs describing them), had significantly better matches between the LLM and neural data than the less commonly linguistically modified subjects (rank sum test z=4.3, p<0.001 **Figure 3J**).

As noted above, there is likely nothing particularly special about Layer 20. Indeed, we found qualitatively similar results for other layers (**Extended Data Fig. 7**). Other features that we associated with the degree of contextual integration included semantic categories (ANOVA F(6264,7)=7.08, p<0.001), part of speech (ANOVA F(6008,8)=5.96, p<0.001), and position within a sentence (r=0.06, p<0.001, Spearman’s correlation, **Extended Data Fig. 8**), though effects were still significant across the parameter space (p<0.05 for all). However, we did not find sentence meaning to be predictive of the degree of contextualization (p=0.74, ridge regression on GTE embeddings versus shuffled models).

### Results are not Unique to the Hippocampus

In two additional patients we recorded single units in the amygdala (P1: 34 units/7346 words, P2: 22 units/3152 words, **Extended Data Fig. 9A**). We were therefore able to examine whether these effects were a unique property of the hippocampus or perhaps a more general property of temporal neurons. We first looked at the neural encoding of position. We found linear encoding of position at rates (36% of units) and amplitudes (mean significant r=0.05, Spearman’s correlation) similar to that of the hippocampus (32%, mean significant r=0.03, Spearman’s correlation, **Extended Data Fig. 9B**). We also find similar rates of oscillatory cells (57% units significant) with similar coverage of the frequency space (**Extended Data Fig. 9C**). Last, we were able to qualitatively recreate the match between neural and LLM contextualization (compare **Figure 3B** to **Extended Data Fig. 9D**), albeit at lower maximum (r=0.12, Spearman’s correlation) with marginals that peaked at an earlier layer (Layer 4) and a smaller optimal window (3).

### Contextual models capture neural variance better than non-contextual models

The fact that word-evoked neural responses depend on preceding words implies that the neural data should be better fit by contextual semantic models than by non-contextual ones^71^. To test this idea, we next compared model fits for contextual and non-contextual embedders (GloVe and Word2Vec), using principal component analysis (PCA) on model embeddings to ensure the same number (100) of predictive dimensions. For each source of semantic embeddings, we created an explanatory Poisson model using ridge regression and compared the log likelihood of the model to that of a model that had its embedding vectors shuffled (ΔLLH). As predicted, the contextual models we tested (GPT-2, Llama-3 and DeBERTa V3, all at layers that maximized the fit to the neural APG) have better fits than non-contextual models (Glove and Word2Vec, sign rank test, p<0.001 for all pairwise comparisons, **Figure 4A**).

**Figure 4.**
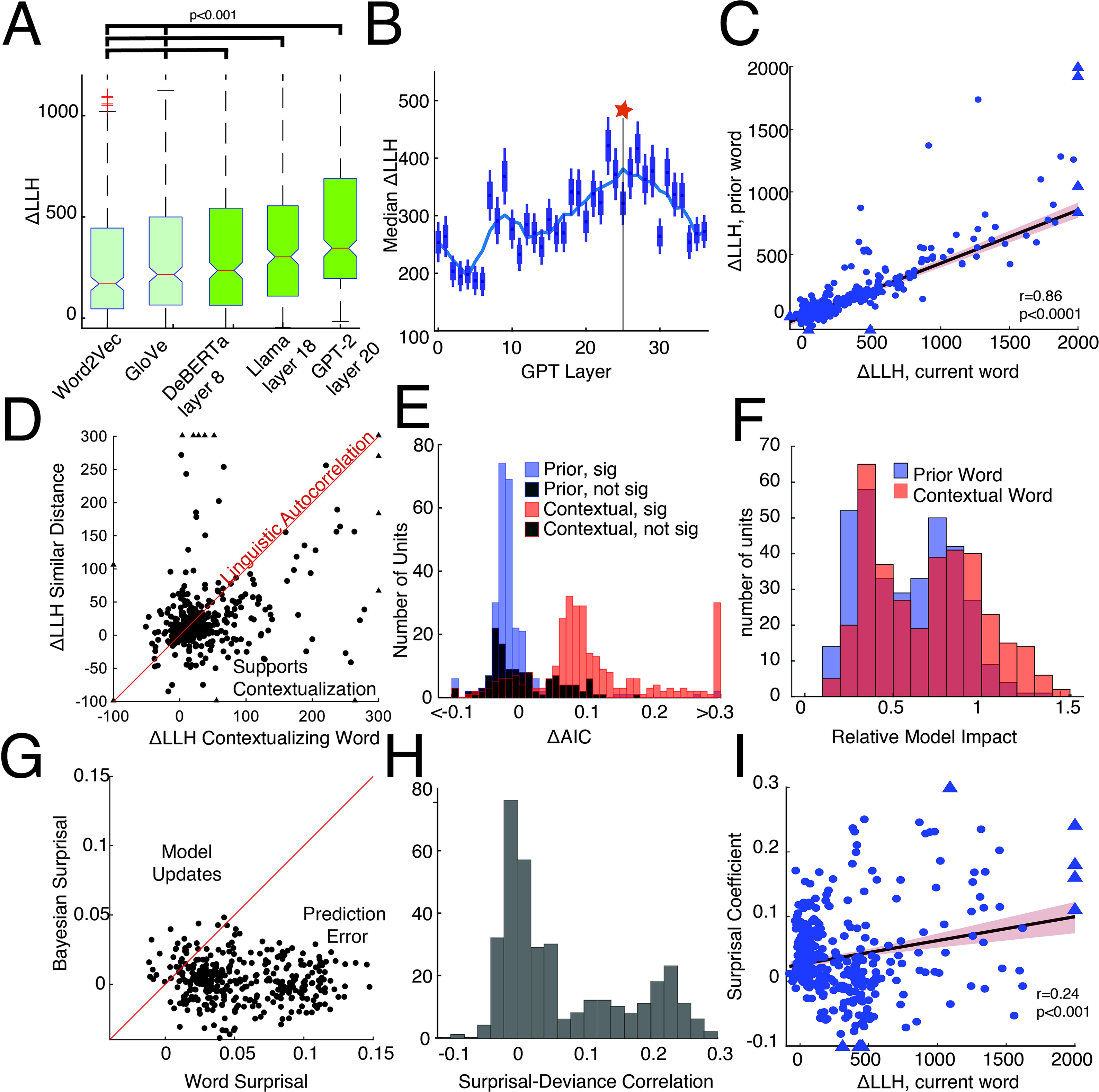
Neural activity reflects combinations of the semantic representations of the current, prior, and expected words. **A.** A comparison of the LLH for each neuronal model (n=365 units) with different semantic embeddings for non-contextual (Word2Vec and GloVe) versus contextual (DeBerta V3, Llama-3 and GPT-2). Boxes are centered at the median and bounded by the 25^th^-75^th^ percentiles. Whiskers are non outlier extrema, plusses are outliers (1.5 times the interquartile range past the first or third quartiles). Wilcoxon sign rank test, p<0.001 for all pairwise comparisons. **B**. Box and whisker (median and 2 times the) plots of the distribution of LLH differences across all neurons (n=365 units) using the hidden states of each layer (n=37 layers) of GPT-2 as the semantic predictors. Blue line represents the median smoothed with a gaussian of width 5 and star indicates maximum of fitted line. **C.** Plot of the Spearman’s correlation of the ΔLLH for current word model versus the ΔLLH for the next word model for each neuron. Triangles are datapoints that are outside the limits of the plot. Black and red shaded lines represent the best fit +\-95% confidence interval. **D**. Plot of the model fits (as defined by the difference in LLH) for models that use contextualizing words versus models that use prior words with a similar distance distribution. The diagonal line represents the hypothesis that there is no difference, consistent with a general effect of autocorrelation. E. Histogram of the difference in the AIC per word between the basic current word model and the current + prior word model (blue) or the current + contextual word model (red). Positive values indicate that the unit’s firing patterns are better explained with both words, even when penalizing for the increased number of parameters. F. Distribution of the norm of the coefficients for the prior word (blue) and the contextual word (red) normalized by the norm of the coefficients for the current word from the joint models, for each unit. G. Plot of Spearman correlations for each unit to surprisal versus Bayesian surprisal. The diagonal line represents a similar impact. H. Distribution of correlations between the deviance of the model for each word and the relative surprise of that word for all neurons for all units. **I.** The ΔLLH of the non-contextual model as a function of the coefficients of surprisal in the linear model that predicts deviance. R value is the Spearman’s correlation. Black and red lines represent the best fit +\-95% confidence interval.

Within a given LLM model, however, contextualization depends on the layer within the model. Typically, higher layers have greater contextualization, although the highest layers may be more oriented towards prediction of potential future words, making them less well correlated with the current semantically turned neural activity^23,72–74^. Consistent with these past studies, we found the strongest model fits at middle layers - in our case, at Layer 25 out of the 37 total layers (**Figure 4B**), which is close to the Layer 20 that optimized the match between neural and LLM APG (**Figure 3C**).

These analyses depend on interpreting the intricacies of LLMs, which include far more than local context. To isolate the effects of context, we next considered the explanatory power of recent words within the current neural representations using a non-contextual embedder (word2vec). Here, rather than quantifying the impact of representations of prior words on those of current words (**Figure 2,3**), we directly examined the relationship between firing patterns and the semantic information of prior words by regressing the neural response to each word against the embeddings of the prior word. We first found a significant relationship in 63.2% of units, though at a lower strength than current words (median ratio of ΔLLH = 40.8 +/- 21.6%). Notably, the same units that responded to the current word also responded to prior words (correlation of the ΔLLH per unit r=0.86, p<0.0001, Spearman’s correlation), demonstrating that the firing patterns of semantically responsive neurons are modulated by both the current and previous words (**Figure 4C**).

Yet perhaps these relationships are due to semantic coherence inherent within autocorrelations in language, in which adjacent words contain similar semantic features. We therefore used the empirical APG derived from GPT-2 to select the most contextually relevant prior word for each word (as per **Figure 3C**). The model using these contextual words to predict current neural firing patterns outperforms a model that picks random words with a similar distribution of word distances (**Figure 4D**, formed by randomizing the distances, Wilcoxon signed rank test on ΔLLH z=4.6, p<0.001). To quantify the added information, we calculated the Akaike information criterion (AIC) of the model using both the current and most contextualizing previous word and compared it to the AIC of the model using just the current word for each neuron. If the two words provide the same information, then this should not improve the model fit. In contrast, we found that adding the immediately prior word improves the fit in 22.1% of the 272 significant units using just the current word (i.e. there is a decrease in the AIC). In stark contrast, adding the LLM APG defined contextual word improved the fit in nearly all, at 94.9% of units (**Figure 4E**) despite being farther away (an average of 2.14 +/-1.3 words prior relative to 1 word prior). Additionally, the representations of the contextual words had a more significant role in the joint models: the norm of the coefficients for the contextualizing words was 70.1%+/-31.5 of the norm for the current words’ coefficients, significantly greater than the 58.8% +/-27.7 for the immediately preceding words (paired t-test, t(355)=13.4 p<0.001, **Figure 4F**). Together, these two results demonstrate that the words used by an LLM to provide context are the best source of information to improve a model fitting ongoing words to neural data. Defining the contextual integration of a neuron as the degree of fit improvement, we found no impact of the units’ location on the anterior-posterior axis (ranksum test z=0.9, p=0.4) or laterality (ranksum test z=1.7, p=0.09).

### Role of context in ongoing next-word prediction

LLMs are trained to optimize next-token prediction, with contextualization an emergent response driven by this objective. We therefore examined the relationship between neural contextualization and next-word prediction in our data. We defined prediction errors as word surprisal, the negative logarithm of the conditional probability of a word given its preceding context. We found a significant correlation between word surprisal and neural firing rates^75^ (mean r=0.06 +/-.002, Spearman’s correlation). In contrast, Bayesian surprisal, which quantifies the difference between prior and posterior distributions over latent representations induced by word presentation^76,77^, explained significantly less variance in neural firing (paired t-test t(355)=24.6, p < 0.001, **Figure 4G**). These findings suggest that neural surprise responses more closely reflect lexical prediction error than signals of model updates.

This model of contextual integration has a testable implication: prediction errors should reduce the faithfulness of neural semantic encoding as they become decoupled from the predictive framework that normally supports neural representation^14^. We defined the *representational error* as the deviance of the models’ prediction for that word. We find an average correlation of 0.07 +/-0.09 between the prediction error and the representational error across units (**Figure 4H**). Overall, 52.0% of units (185/356) show a significant positive correlation versus just 8.2% that show a significant negative correlation (29/356) (α =0.05). In other words, surprising words are represented less faithfully than unsurprising words. Nonlinear neural dynamics elicited by high-surprisal words could also plausibly contribute to the reduced model fit we observe for these events. However, this account likewise implies that surprising words engage neural processes that deviate from baseline coding states. Thus, regardless of model form, the data indicate that hippocampal neurons are sensitive to deviations from expected word distributions, expectations that must be constructed from the preceding context.

Artifactual explanations for this result could include that surprising words are less common, can occur in moments of uncertainty (high entropy), or even just elicit a higher firing rate (any of which could decrease the accuracy of representation). We can control for these possibilities by creating a linear model that predicts the deviance of each word using the surprisal, frequency, entropy, spike count of the word, and all of their interactions. Doing so, we nonetheless still found a significant positive effect of surprisal in 32.9% of units (versus a negative effect in 2.5% of units). Interestingly, despite the significant relationship between the accuracy of the model in representing the current word and the impact of surprisal on its deviance (r=0.24, p<0.001 Pearson’s correlation), it was not always the units with the highest semantic accuracy that had the strongest relationship between model error and surprisal (**Figure 4I**). This weak correlation at the level of units implies there is an unavoidable tradeoff between accurate representation of the current word and susceptibility to modulation by ongoing predictions.

To test the robustness of our findings, we repeated our analyses using an alternative pipeline that (a) generated null distributions via circular shifts instead of random shuffling^78^, (b) assessed model performance on held out data with 5-fold cross validation, using a predictive rather than explanatory framework^79^. Doing so, we grossly find qualitatively matched results (**Extended Data Fig. 10**). Specifically, contextual models capture neural variance better than non-contextual models (**Extended Data Fig. 10A**, sign rank test p<0.001 for all pairwise comparisons), Layer 23 optimizes the fit (**Extended Data Fig. 10B**), units responded similarly to prior words (**Extended Data Fig. 10C** r=0.61, p<0.0001, Spearman’s correlation), adding contextual words improved the AIC in 85.4% of units (albeit only when using the same size design matrix, see Methods), prediction error and the representational error were correlated (mean r= 0.07 +/-0.09, Spearman’s correlation), and units with higher encoding accuracy still showed a stronger impact of surprisal (r=0.51, p<0.001 Spearman’s correlation). Contextualizing words fell below significance (**Extended Data Fig. 10D**), however, likely due to decreasing the testing dataset size by a factor of 5.

## DISCUSSION

We find that hippocampal representations of word meaning incorporate the activation patterns associated with the words that contextualize them. Further, the degree of this incorporation is similar to attention weights used by LLMs. These results therefore suggest a specific mechanism for semantic contextualization in the human brain, one that relies on averaging of patterns associated with contextualizing words.

Neural codes for positions of words provide key information for contextualization, as they represent a mapping principle^80–82^. This result, then aligns with the broader mapping functions of the hippocampus which include space, time, and concepts^83–85^. The finding of a second, independent sinusoidal positional encoding is particularly interesting. Periodic codes have benefits including representation of relative distances, smoothness, generalization to arbitrarily long sequences, and ease of learning (relative to learned positional representations^86^). These factors have also been proposed for why the medial entorhinal cortex has sinusoidal codes for navigational space in addition to hippocampal place cells, in the form of grid-cells^85,87^. Our results, suggest the extension of the principle of periodic codes to the domain of language

Note that position information does not need to dominate the embedding space-in our data for example, ramp coding could explain only 1.5% of neural variance. Similarly, in many LLMs dimensions associated with absolute or relative position appear to occupy a comparatively small, specialized subspace of the overall representation, rather than dominating the embedding geometry^88^. We emphasize that the bias towards later words in a sentence is unlikely to reflect a growing summation of stimulus evoked responses. Our linguistic stimulus is a continuous natural stimulus containing a series of words which rarely had pauses between words or sentences. Further evidence that these results do not reflect a stimulus-evoked response comes from the observation that the growth is observed at multiple scales (phrase, sentence). Last, this response is greatly reduced in Jabberwocky stimuli, which have similar acoustic and syntactic information but are lacking semantic information, suggesting that positional tracking is dependent on the semantic content.

Not all word pairings show equal attentional modification, raising the question of what drives the difference. For example, subject nouns did not strongly modify their subsequent verbs, whereas verbs did strongly modify their subsequent object nouns. (This is a particularly clean example, because it is a strict reversal,). Notably, this pattern was observed in both neural and LLM representations, suggesting it is a general feature of language (**Extended Data Fig. 7**). This observed shift following well-established linguistic principles, which in this case can be explained in information-theoretic terms: the word “person” can have any of a large number of verbs after it, whereas a verb like “swim” or “eat” or “sing” can only refer to nouns capable of these actions. So the verb therefore has a larger constraining effect (and thus attentional weight) on its object noun, whereas the subject noun has a more modest constraining effect (and lower attentional weight) on its verb.

Contextualization in the brain does not happen arbitrarily; instead, it is a learned process that attempts to achieve a specific goal. Understanding this “objective function” is a key question in linguistics. Computer scientists have independently found that next-word-prediction (or, more generally, next-token prediction, see below) is a tool powerful enough to introduce features not only remarkably similar to contextualization, but also other cognitive processes such as syntactic parsing^89^, reasoning, and generalization^10,90,91^, though see ref. 92. In other words, many of the core features of language are not built into LLMs; instead, they are an emergent feature of architecture and training principles. Our results here highlight the links between contextualized responses and next-word prediction in the human brain. The relationship between neural activity and upcoming words has been interpreted as preparatory activity^90^, though others have raised concerns that this analysis is limited by inherent correlations within natural language^93^. We therefore focused on prediction error, demonstrating that information about upcoming words must be present, as the evoked responses were modulated by these predictions. Further, we directly tie prediction errors to semantic representations, showing that surprise causes more than an error signal-it changes the neural response to a word. Future research could investigate how prediction errors modulate learning signals in the hippocampus, and how these signals could contribute to the refinement of semantic representations.

The hippocampus is well-situated to perform the core elements of semantic contextualization: (1) it is closely associated with mapping functions that are related to temporal position encoding^94,95^, (2) it integrates information across multiple modalities and uses that information to contextualize representations in a variety of domains^96^, (3) and it is closely associated with the processes of prediction that drive contextualization^97,98^. Finally, (4) the hippocampus has a long-established role in memory, one that has many links with semantic knowledge and with semantic representations in particular^37,99^. Indeed, the role of the hippocampus in language is well established^43,44,46^, as is its role in semantic representations specifically^32^. Recent studies using intracranial unit recordings have begun to show clear evidence for hippocampal role in language^28,32,75^. We suspect one reason for the absence of the hippocampus in classical language models is negative evidence - the hippocampus can be difficult to image using fMRI and lesions isolated to the dominant hippocampus are rare. However, another factor is that the hippocampus is clearly not a site uniquely aligned to language coding processes, so it does not fit modern uniqueness-focused definitions of language areas^8^.

### Limitations

Our study has several limitations. First, by having only one “shift” vector per word we can only examine the potential neural correlates of the sum of the attentional weightings. This limits us from examining correlates of individual attention heads which may specialize for different grammatical and semantic roles^100^. This simplification likely overlooks a dynamic and distributed attentional mechanism in the brain, wherein different heads may attend to different features^12^. Second, we ignore the fascinating multi-layer perceptron (MLP) element of contextualization^16^. This nonlinear step accounts for more model parameters than the attention block and allows for many higher order features, such as representation that distinguishes learned (rather than inferred) contextualization^101^. Additionally, several of our results are quite low in absolute magnitude. This is expected given that we are performing, in essence, single word (or word-pair) analyses, rather than averaging over multiple trials. On the other hand, we have responses of hundreds of neurons, giving us strong sensitivity even with low correlation values. This pattern is observed in other studies that have low numbers of repeats but large datasets^33,45,102–104^. Moreover, our results are almost entirely restricted to the hippocampus. Notably, we were able to replicate core findings in a subpopulation of amygdala recordings. Perhaps, however, this should not be surprising, as in general conceptual representations are shared in the hippocampus, amygdala, and indeed other medial temporal lobe (MTL) regions^105^. In any case, it would be particularly interesting to consider the computational mechanisms of contextualization in language regions in the lateral cortex.

Both language and neural data have a natural temporal autocorrelation structure that may cause spurious positive effects in studies such as this one. This autocorrelation should cause a clear decay curve in self-similarity measures for both neural and linguistic data. Indeed, we find a clear temporal autocorrelation structure in our own data (Figure 2C). What is important, then, is the ways in which our data go beyond this autocorrelation structure - the fact that deviations from the decay curve match in both neural and APG data (e.g. Figure 3A,D). Further, while activity in hippocampal neurons does reflect prior words, we show that the contextualizing words (which can be a few words away) have the largest impact (e.g. Figure 4EF).

Finally, another limitation of our findings is that our results are limited to English speakers. English has weak morphology and strict syntax, compared to languages like Warlpiri, Kalaallisut, and Latin, which have freer word order. Would word order be represented in the same way in these languages? On one hand, the effects we attribute here to word order may, in reality, reflect syntactic roles, in which case responses would follow words as they move around. On the other hand, apparent word order effects may reflect a core word order encoding, used to scaffold meaning, and may stay the same regardless of word order. These debates in turn have parallels in debates about the nature of place coding in hippocampus, and, in particular, whether it reflects a robust place scaffold or behaviorally relevant maps^95,106–108^.

## METHODS

### Experimental Paradigm

We recorded single-neuron activity from the hippocampus of ten adult patients with drug-resistant epilepsy (six male, four female), age range 18 to 45 years old (mean 29.1 +/-8.1), undergoing intracranial monitoring at Baylor St. Luke’s Hospital for seizure localization. All English speaking patients diagnosed with drug-resistant temporal lobe epilepsy that were scheduled to undergo intracranial monitoring for the duration of our study were offered a chance to participate. All patients gave informed consent. Patients were compensated at standard rates as approved by our IRB. None had hippocampal seizure foci. Electrode position was confirmed on postoperative CT^109^. We used Behnke-Fried stereoelectroencephalography (sEEG) probes and a Blackrock Neuroport system. Spike sorting was performed with WaveClus^110^ and carefully sorted offline by highly trained lab members using a standardized rubric^32^. Participants listened to six Podcasts from *The Moth Radio Hour* (47 minutes; 7,346 words). Audio was synchronized with neural recordings and then transcribed by AssemblyAI and corrected manually using PRAAT^111^.

### Generation of Word Embeddings

We used three word level LLMs: GPT-2 Large^112^, Llama-3^69^, and DeBERTa V3^68^.

#### GPT-2 Large

We extracted 37 hidden states consisting of 36 transformer blocks + 1 embedding layer. Each word was a 1280-d vector, and the native context window was 1024 tokens.

#### Llama-3 7B

We extracted 33 hidden states consisting of 32 transformer blocks + 1 embedding layer. Each word was a 4096-d, vector and the native context window was 1024 tokens. For both GPT2 and Llama-3, we built an expanding context window. Beginning with an empty string, we appended one new word, re-tokenized the running text, ran the model in inference-only mode, and averaged sub-word vectors belonging to the just-added word. After 1024 tokens, we truncated the leftmost words so that all three models would be compared under identical context windows.

#### DeBERTa-V3

Base (12 transformer blocks + 1 embedding layer, 768-d vectors, 512-token context)^68^ was handled separately due to its bidirectional encoder architecture, which processes the entire input in parallel. To align outputs with the unidirectional processing of GPT-2 Large and Llama-3, we applied the expanding-context procedure described above. Although the model sees both past and future tokens, we extracted only the hidden states corresponding to the just-added word which limits interpretation to left-contextual information. Sub-word vectors were averaged to yield one vector per layer. Hidden states from all layers were retained.

For a separate analysis, we used one sentence level LLM.

#### GTE-large-en-v1.5

(24 transformer blocks + 1 embedding layer, 1024-d vectors, 8192-token context) is a bidirectional encoder trained with a large-scale contrastive retrieval objective^60^. Unlike the progressive word-level extraction used for decoder-only models, each full sentence was tokenized and processed simultaneously. Hidden states from all layers (embedding layer + 24 transformer blocks) were retained. For each layer, sentence-level representations were computed using attention-masked mean pooling across token embeddings. This yielded one 1024-dimensional vector per sentence per layer (25 total layers). Only the final layer was used for analysis. Because GTE processes each sentence bidirectionally and integrates information across all tokens within the segment, embeddings reflect holistic thematic representations that incorporate the contextual structure preceding the sentence.

We used Word2Vec and GloVe to derive static embeddings. All words were converted to lowercase and stripped of punctuation before lookup. Surnames and proper nouns that were not present in the embedding vocabulary (e.g., “*Applebee’s*”) were excluded from downstream analyses. For Word2Vec we employed the pre-trained fastText skip-gram model^113^. For GloVe, we used the Common Crawl model^114^ (glove.840B.300d.txt).

### Neural Analyses

Neural responses were defined as the spike count during the word presentation, shifted by an offset of 80 milliseconds to account for response latency.

### Ramp Coding

Sentence boundaries were generated by AssemblyAI’s punctuation-restoration model and confirmed by a trained annotator. Clause boundaries were initially suggested by automated Penn-Treebank–style parses (CoreNLP^59^) and then manually adjusted by a trained annotator to match standard subordinate-clause structure and to correct any parser errors.

For each unit we defined ramp coding as a significant correlation between firing rate and word index within a sentence. To look at the impact of the GTE sentence level embeddings on ramp coding of position we reduced its dimensionality via PCA to the first 10 components, and included position, these components, and the interaction terms as regressors in a linear model relating these values to firing rate.

### Index Coding

For each hierarchical level, we performed a Kruskal-Wallis test on firing rate responses to each word, grouping by index up to the first 10 indices (to maximize sample number). Cells that demonstrated a significant main effect were further analyzed with pairwise rank-sum tests. Significant differences were visualized using heatmaps, with cells sorted by the first group showing a significant increase. To assess whether single-index selectivity exceeded chance, a binomial model was used assuming a uniform probability of 0.1.

### Fourier coding

We used Welch’s power spectral density (PSD) estimate (range: 100 frequencies between 0.05 and 0.5 Hz). We used windows of 512 words (overlap: 256 words). Rare (30/7356 words in 1/10 patients) missing words from experimental interruptions were linearly interpolated between adjacent values. PSDs were smoothed with a gaussian filter that had a full width half maximum at 0.023 Hz (15 samples). We estimated supra-aperiodic peaks using the FOOOF toolbox^65^ and selected the largest peak. Units that did not have a peak with a minimum width of 0.01 Hz were considered non oscillatory. Statistical significance was defined with a double threshold: (1) Noise was defined as the residual after removing the aperiodic fit. Significant peaks needed to be at least 2.5 standard deviations of the noise estimate above the aperiodic fit. (2) Firing rates were shuffled 100 times per unit and the same analysis pipeline was applied. Peak amplitudes for each unit from each shuffle were recorded. Significant peaks needed to be larger than the 95th percentile of this distribution.

We also calculated the intrinsic frequency of the units during the resting period, defined as the time before and after the experiment. Spike count was sampled at 1 kHz. Here we used windows of 4.4 minutes and 2.2 minutes of overlap to obtain similar frequency resolution. The frequency range was 0.045 to 4.5 Hz, corresponding to a median word rate of 4.5 words per second. For example, if a neuron fired once every other word, this would correspond to a preferred frequency of approximately 2.3 Hz.

### Attention Pattern Grid

The ***neural attention pattern grid*** is an estimate of the degree to which all past words influence each current word. It recapitulates the function performed by the APG used in transformer architecture LLMs. We first estimated the neural APG from neural data alone (no LLMs) and then, for comparison, performed an analogous process on the LLM embeddings. Note that the LLM-derived empirical APGs were not derived directly from the LLM APGs themselves; they were inferred so as to keep the estimation procedure the same for brain and LLM. Embedding matrices were 7,346 rows, corresponding to the number of words in our stimulus set, by M columns, corresponding to the number of embedding dimensions per word. For the neural embedding matrix M was equal to 356, corresponding to the number of neurons collected across patients. For the semantic empirical APGs, M ranged from 300 (word2vec) to 4,096 (Llama-3).

We then created two new embedding vectors for each word: the average vector across all instances of that word 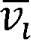 and the shifted representation of the word j is 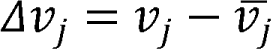 Then each value of the APG was created by measuring the cosine similarity between the shifted vector and the mean vector of the previous words

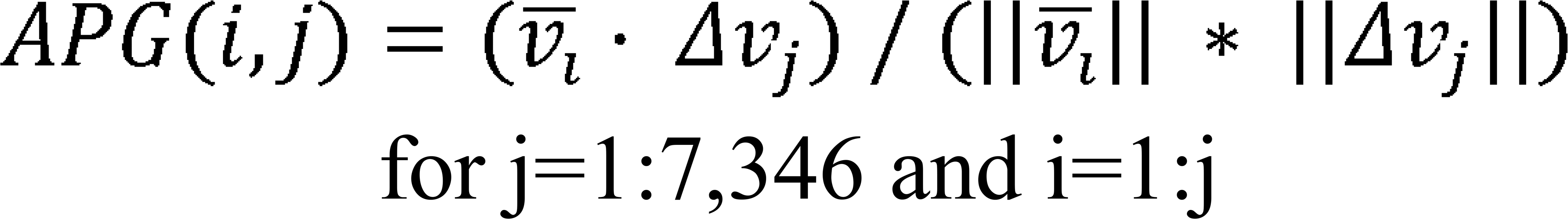

Words with only one instance in the stimulus set were excluded from analysis as it was impossible to calculate a shifted representation. Only causal influences were calculated.

### Clustering Attention Vectors

To explore the structure of our empirical APGs, we constructed a k-nearest neighbors (k-NN) graph on truncated APGs going back 5 words. This graph served as the basis for unsupervised clustering. We applied the Leiden algorithm to segment the graph into distinct groups of words with similar recent attention dynamics by maximizing within-pattern similarity while minimizing similarity between patterns in the APG similarity graph. These clusters were hypothesized to reflect shared structure patterns in word relationships.

### Assessing the impact of sentence meaning on contextual integration

We regressed the top 100 PCA dimensions of the sentence embeddings against the match between neural and LLM APGs. All words in the same sentence had the same embedding. The R^2^ value was defined as the median R^2^ value tested on the held-out data across folds, and the maximal value was taken across lambdas. For significance testing words were circularly shifted by a random number and the regression was run 1000 times and the resultant R^2^ distribution was used to define chance.

### Poisson Regression

Word-evoked neural responses were defined as the number of spikes occurring for the duration of that word, with an offset of 80 milliseconds. Each word was also represented by a feature vector (e.g., GPT-derived semantic embeddings). For each feature, PCA was used to generate the 100 orthogonal dimensions that explained the most variance. If two features were used (e.g. current and prior word) then they were concatenated. A scalar **duration** vector represented the duration of each word. Each column of **X** was standardized to have zero mean and unit variance after being split into folds (see below) to avoid data leakage. Additionally, 100 shuffled versions of **X** were generated by permuting the rows of **X_raw**, recomputing the interaction terms, and standardizing again. These permutations were used to generate null distributions for hypothesis testing. Durations were not shuffled.

#### Poisson Ridge Regression Model

We modeled spike counts with a Poisson generalized linear model (GLM) using an exponential link function. For each word *w*:

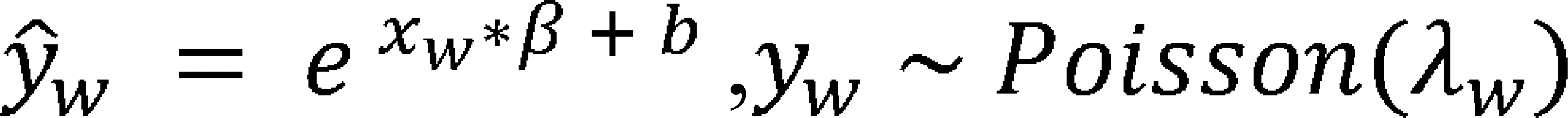

Where:.*x_w_* is the feature vector for word w, is the weight vector, *b* is a scalar bias, and is the predicted spike count, and *y_w_* is the observed spike count.

We used ridge regression to prevent overfitting, yielding the following loss function:

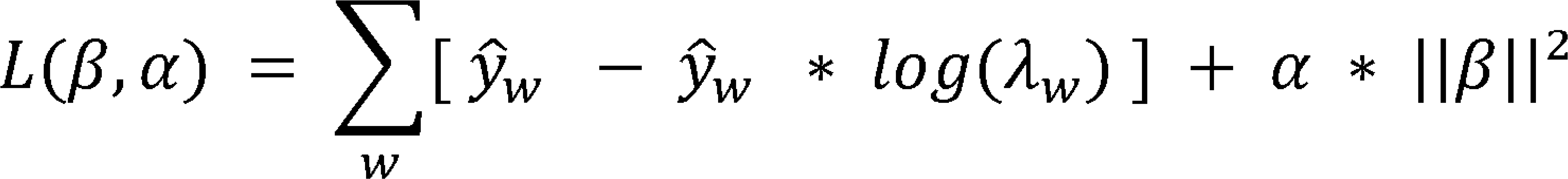

wherein _α_ is the regularization parameter.

The model was implemented in PyTorch on a NVIDIA GeForce RTX 4070 Ti SUPER to decrease training time. Training was performed using the Adam optimizer^115^ with a default learning rate of 0.01. Training proceeded for up to 300 epochs, with early stopping if the decrease in penalized loss fell below 10^−4^ for ten consecutive epochs. Loss was computed at each training epoch. Gradients were back propagated, and the weights were updated until convergence or early stopping.

#### Hyperparameter Selection

To choose the regularization strength α, we used a two-stage grid search:

1. **Coarse Search** over: 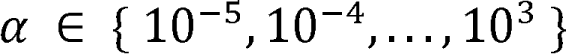
2. **Fine Search** around the best coarse value, spanning two orders of magnitude.

For each value of _α_, we performed 5-fold cross-validation. Folds were temporally contiguous to avoid data leakage. For each fold, the model was trained on the other four folds and evaluated on the held out fold by computing the Poisson log □ likelihood:

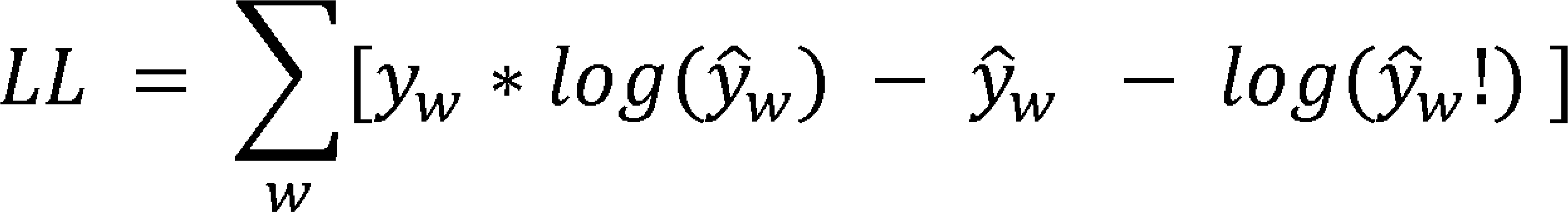

These fold wise log likelihoods were averaged to yield an overall score for each _α_. The _α_ yielding the highest average log-likelihood was selected.

#### Final Model Fitting and Evaluation

Once the best α was determined using cross-validation, we created an explanatory model by training it on all available data for each neuron using the selected α.

We computed the following evaluation metrics:

● Log-likelihood of the model (as above)

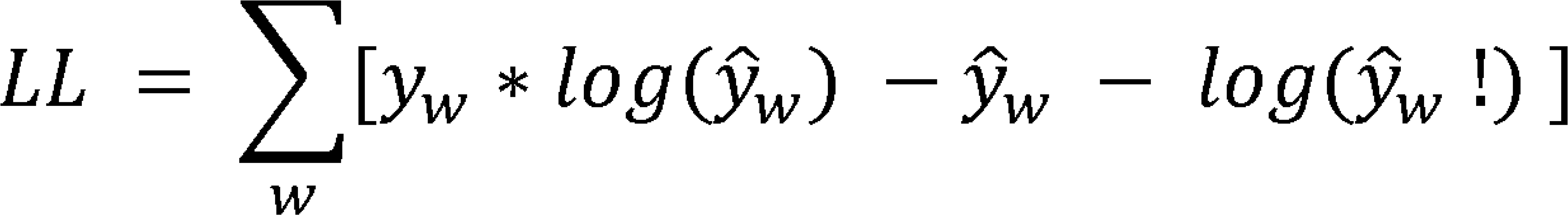

● Deviance of the residuals for each word:

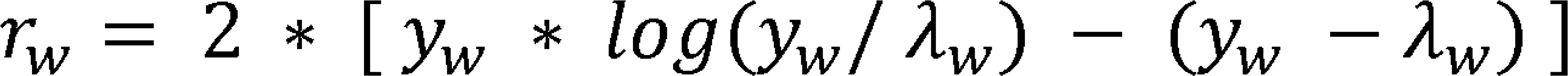

#### Permutation Testing

To determine whether the observed model performance exceeded what would be expected by chance, we computed a null distribution of log-likelihoods using 100 permuted feature matrices. For each permutation:

1. A Poisson ridge model was fitted using optimal _α_ determined by the true data.
2. Log-likelihood was computed. The empirical p-value was defined as

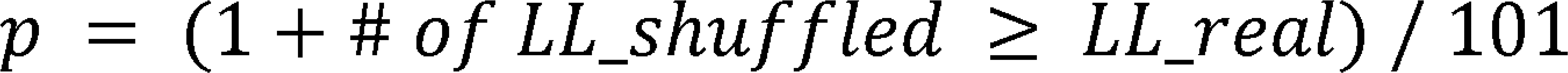

#### Alternative Model Decisions

To evaluate robustness we repeated our analyses using alternative design decisions. Alternative models were trained using a ridge-regularized Poisson generalized linear model (GLMs) and evaluated using 5-fold outer cross-validation. Within each fold, regularization strength was selected to maximize a 5-fold inner cross-validation. All design matrices were the same size (100 PCA dimensions). For models that tested combinations of words (e.g. current word + contextual word), every other PCA dimension was used from both (e.g. current word would be odd dimensions and contextual word would be even dimensions) to create equity between the models. For each neuron, metrics described above were calculated on the held out data for each fold and aggregated using the median across folds to reduce sensitivity to outliers. Deviance was calculated on each fold and aggregated across folds. To assess significance, we generated 100 shuffled predictor matrices and circularly shifted the predictor matrix along the observation axis by a randomly selected offset of at least ±100 samples (to avoid short-range autocorrelation artifacts).

### Surprisal and Entropy

For each word we slid a progressive 1,024-token window through the preceding transcript and fed this context into *GPT-2-large*, and extracted the probability distribution over the vocabulary for the current word. From this distribution, we computed two standard information-theoretic measures:

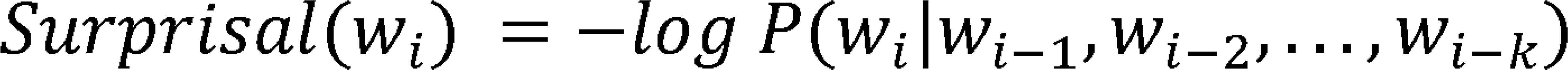

the unexpectedness of the actual word *w_i_*, and

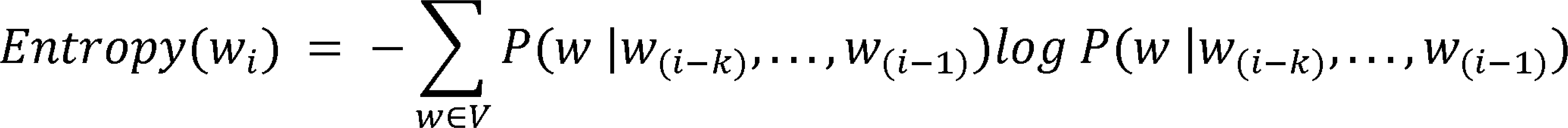

which quantifies the model’s uncertainty over all possible next words in the vocabulary.

### Dependency Grammar Relations

We derived governor–dependent pairs from each utterance using the Stanford CoreNLP dependency parser^59^. After reconstructing sentences from the transcript, every sentence was sent to a locally hosted CoreNLP server where it was parsed on a locally hosted CoreNLP server; the parser outputs the directed, labeled relations of the Stanford Typed Dependencies scheme^116^, closely aligned with Universal Dependencies^117^. Because CoreNLP tokenizes contractions and multi-word expressions (e.g., “t”,“-”, “shirt”) differently from the manual transcript, we implemented a post-hoc merging routine: special three- and two-token patterns and common clitic suffixes were collapsed.

### Part of Speech Pairing & Subject and Object Pairing

We adopted the Stanford CoreNLP POS (part of speech) Tagger, which assigns fine-grained part-of-speech labels as per the standards of NLTK^118^. These granular labels were also grouped into broader categories (e.g., “Noun,” “Verb,” “Adjective,” “Adverb”). Next, for each sentence we queried a local CoreNLP server for enhanced dependency graphs (see **Dependency Grammar Relations**^116^). We built six part of speech-defined governor–dependent relations and one subject–object relations:

● Verb–Noun: verb governs noun via *nsubj, nsubj:pass, obj, dobj,* or *iobj*. We distinguish distinguishing subject-first (“noun ← verb”) from object-after (“verb ← noun”).
● Adjective–Noun: identified through the *amod* label in either direction. We distinguish *pre-nominal* (adjective before noun) from *post-nominal* (adjective after noun).
● Preposition–Noun: identified by *case*, which bind a preposition to the head noun of its prepositional phrase.
● Determiner–Noun: identified by *det* (e.g., *the* → *book*).
● Adverb–Verb: identified by *advmod*.
● Possessive Pronoun–Noun: identified by *nmod:poss* and *poss* (e.g., *her* → *idea*).
● Subject–Object triplets: created only when the same verb simultaneously governed at least one *nsubj* (or *nsubj:pass*) and one *obj/dobj/iobj*.

### Opening Nodes, Closing Nodes, Syntactic depth

Following prior work^57,58^, we treat syntactic depth as a simple count of how many constituent phrases remain open above a word in a Penn-Treebank parse. In practical terms, this equals the number of open brackets at the position of *w* when the tree is read left-to-right. For example, in the sentence “The boy who ate the apple slept,” the word “boy” has few open constituents (Sentence, Noun Phrase), and has low syntactic depth and minimal phrase closure. By contrast, “apple” has several nested constituents, resulting in a higher closing-node count and greater syntactic depth. Thus, syntactic depth reflects how many hierarchical phrases are currently active, while opening and closing nodes index where new constituents begin or existing ones terminate in a sentence.

### Hierarchical Phrase Indexing

We developed a novel measure to capture the nested phrase structure that surrounds every word. We converted each Penn-Treebank parse of a sentence into a depth-by-word index matrix (see **Opening Nodes, Closing Nodes, Syntactic depth**). We store these local indices in a matrix M whose rows correspond to successive depths and whose columns correspond to transcript words. Thus, the first row lists the running word indices, while each deeper row re-indexes words relative to their surrounding phrase at that depth: identical numbers within a row indicate membership in the same NP, VP, CP, etc. Because the *ROOT* placeholder occupies Depth-1, top-level phrases (e.g., the sentence’s main NP and VP) appear at Depth-2, ensuring that deeper levels faithfully reflect increasing syntactic embedding.

### Jabberwocky stimuli

Seven patients listened to “Jabberwocky stories” (*1797* words) generated by “Wuggy”^61^, which alters stimulus words to make them meaningless but preserves syntactic structure by retaining function words and morphemes. For example “Before I smow it, the foodbace toom is at my frose woor, and a coakle of them have bapes of bour under their alk.”

## Data and materials availability

## Acknowledgements

We thank Joshua Adkinson, Hanjie Chen, Justin Fine, Victoria Gates, Suzanne Kemmer, and George Kokalas for invaluable assistance.

## Authors contributions

JB and TI contributed to this work equally

KK JB TI MF EM SP SS and BH conceptualized the analysis.

KK JB TI MF and EM performed the analysis.

DP and SS implanted the leads.

AGC AC MF EM RM EB NP AW SS and BH ran the experiment.

KK JB TI SS and BH wrote the manuscript.

All authors contributed to the conceptualization and final form of the manuscript.

## Ethics statement

Experiments were conducted according to protocol guidelines approved by the Institutional Review Board for Baylor College of Medicine and Affiliated Hospitals, Houston TX (H-18112).

