## Extended Data Figures for "Linguistic contextualization in the human hippocampus"

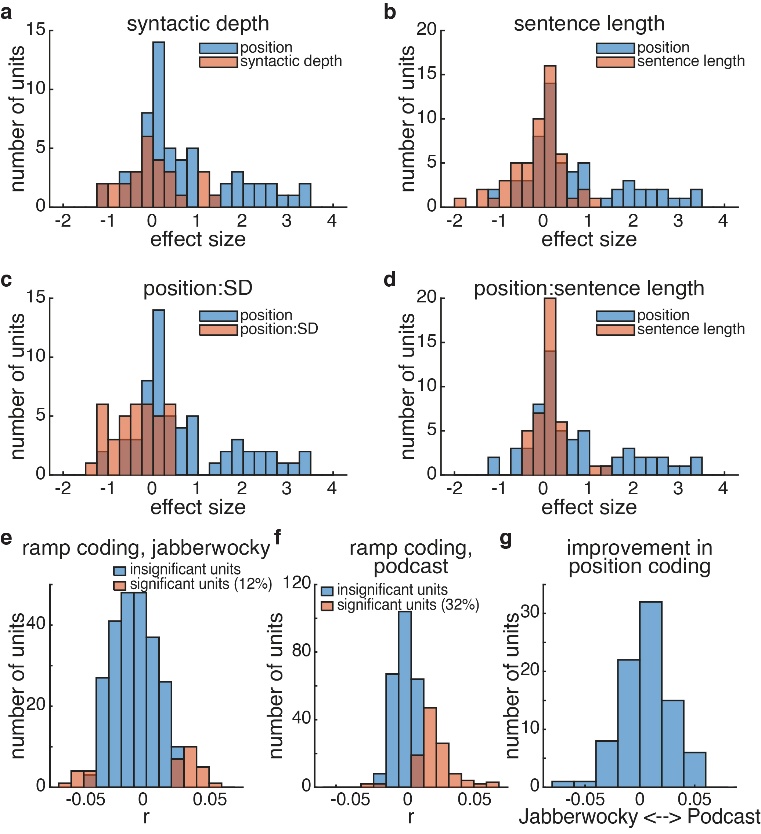

Extended Data Fig. 1: Modulators of ramp coding of position. A. Comparison of the effect sizes of the significant units for position versus that of syntactic depth. B-D. similar histograms but for sentence length, the interaction between position and syntactic depth, and the interaction between position and sentence length, respectively (p<0.01 for all, ranksum test). Ramp coding in Jabberwocky data. E. Distribution of correlations between firing rate and word index in a sentence for all (blue) and significant (red) units in the Jabberwocky Task. F. Similar to E but the podcast task. G. the difference in r values between the podcast task and the Jabberwocky task, showing weak and inconsistent effect

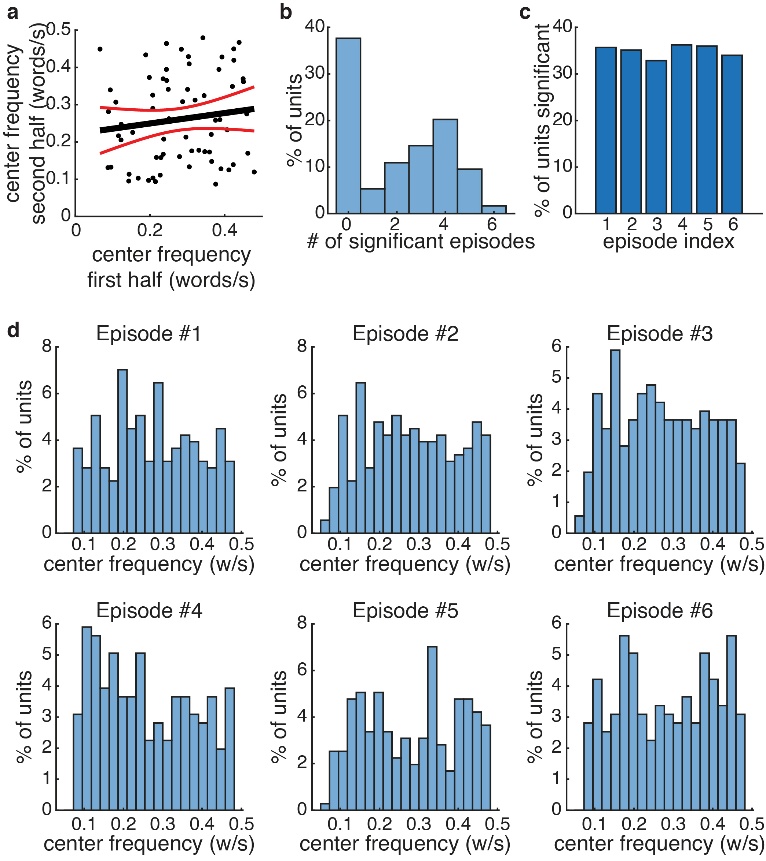

Extended Data Fig. 2. Stability of Unit Identity but Not Center Frequency. A. Center frequencies of units with significant oscillations in first and second half of the dataset. Black shows best fit with red lines showing errors bars of fit, which includes a slope of 0. B. Number of episodes for which each unit had a significant center slope. C. Number of significant units per episode, mean 35.0 +/- 0.5%. D. Distribution of center frequencies of significant units for each episode.

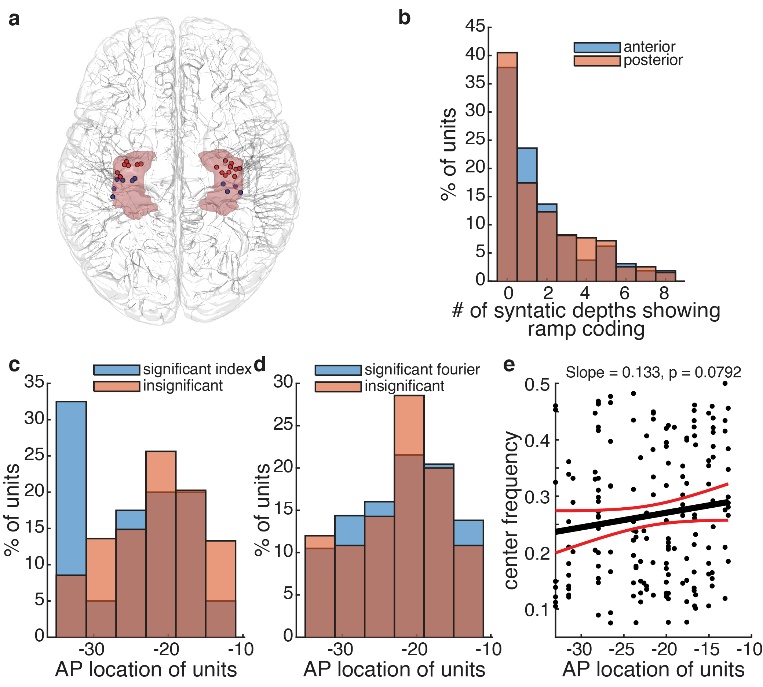

Extended Data Fig. 3: Impact of Anterior-Position location on position coding in the hippocampus. A. Locations of microelectrodes split by anterior (red) and posterior (blue) hippocampus, similar to Figure 1A but from a superior view. B. Histogram showing the percent of units as a function of the number of syntactic depths for which each unit is significant. C. Locations of units significant for Index Coding in MNI coordinates along the anterior posterior axis. Anterior and Posterior are defined at the border of -21. D. Locations of units significant for Fourier Coding in MNI coordinates along the anterior posterior axis. E. Center frequency of units as a function of their anterior-posterior location. Red lines indicate 95% confidence intervals.

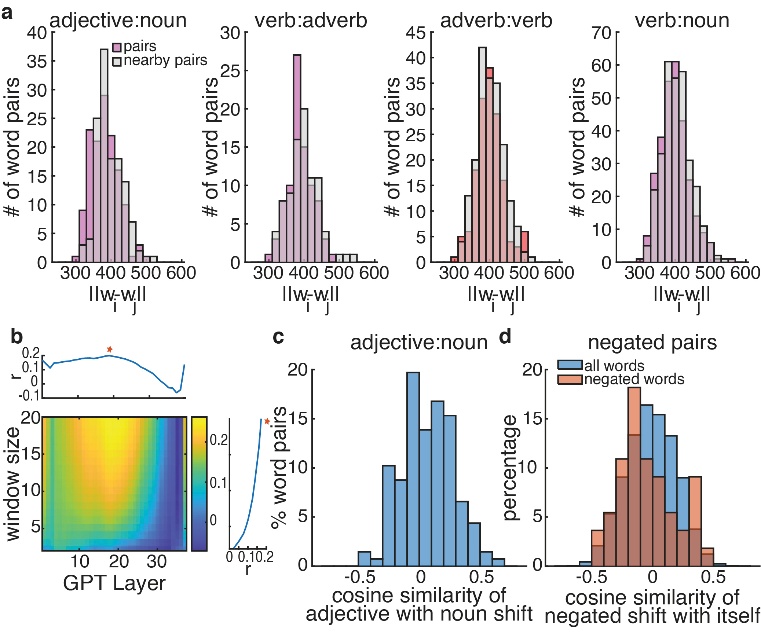

Extended Data Fig. 4. Robustness of findings to distance metric used. A. Reproduction of Figure 2 D-G using Euclidean distance. Note that stronger similarities would be negative here. B. Reproduction of figure 3B using Euclidean distances. C. Distribution of cosine similarities between the vector representations of adjectives and the shift from the mean representations of their respective nouns. D. Distribution of cosine similarities between the vector representations of words and the shift from this mean representation specifically when there is a negative in front of them (e.g. “not know”). Blue distribution represents all words

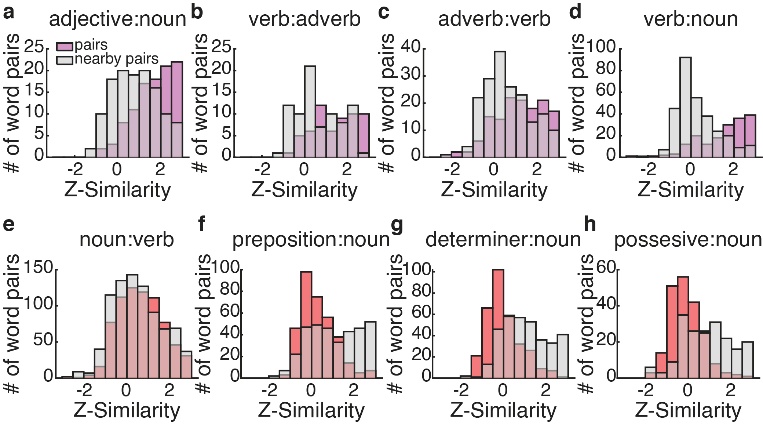

Extended Data Fig. 5. Distributions of word pairs’ contextual strength in GPT2 are similar to neural data. A-H: Distribution of the z-scored contextual pairing values for the word pairs designated above (purple is significant, red is not significant) and the values for control word pairs (gray) using the empiric APG generated from GPT2 layer 20.

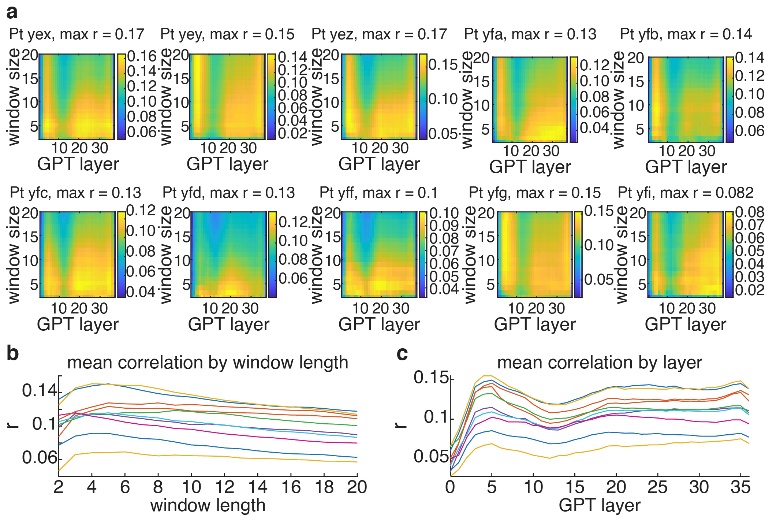

Extended Data Fig. 6: APG analysis can be recreated on an individual patient basis. A. Individual APGs formed from neurons unique to each of the 10 patients. Unique patient ID and maximum correlation value is indicated in the title. B. Marginal distribution for each patient (indicated by different colors) of the correlation as a function of window length. C. Similar to B but as a function of GPT layer.

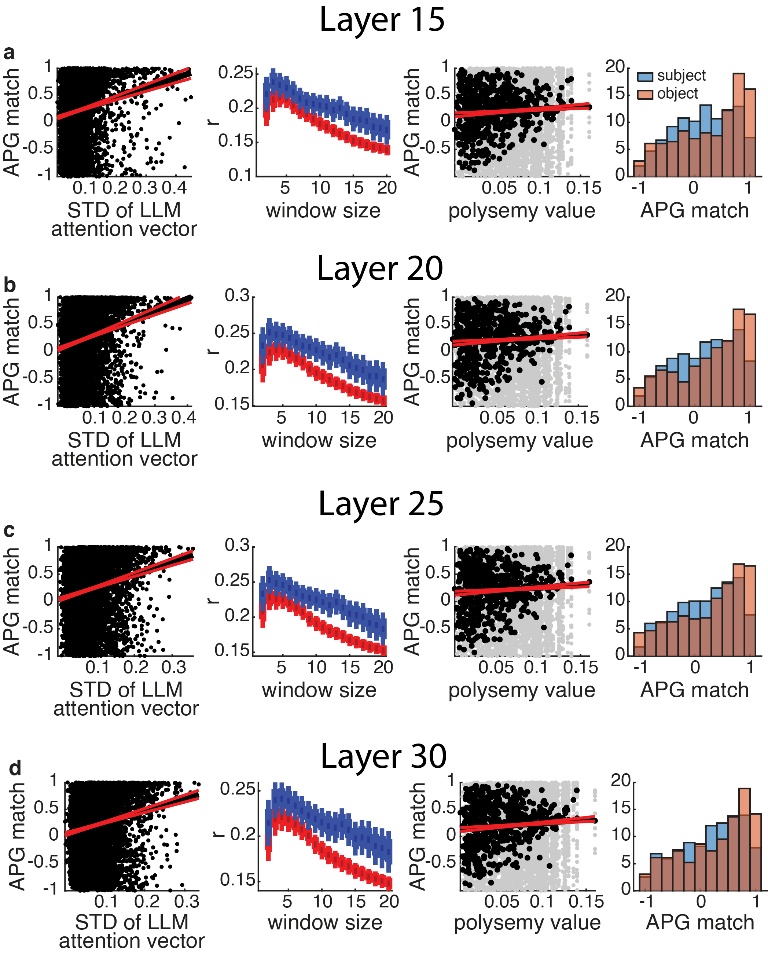

Extended Data Fig. 7 Results do not depend on GPT Layer. Reproductions of Figure 3G-J with GPT Layers 15 (A), 20 (B), 25 (C) and 30 (D). All statistical testing was still significant (p<0.006 for all).

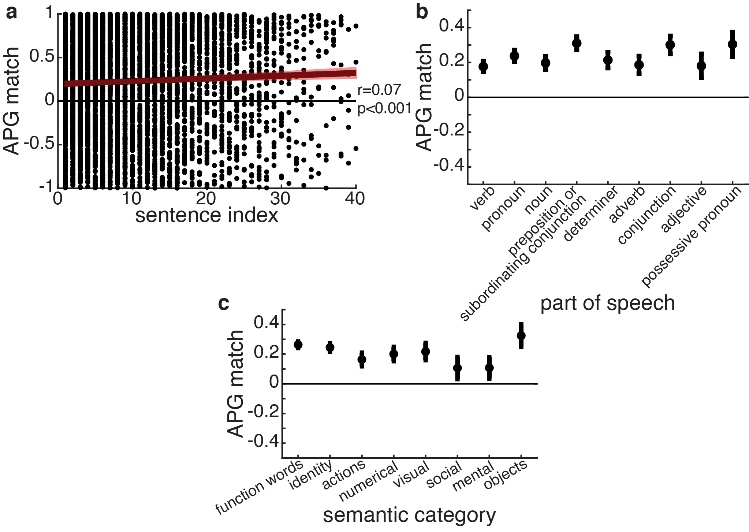

Extended Data Fig. 8: Modulators of APG match. We observe a correspondence between neural and LLM data as a function of word position within a sentence. Red shading shows 95% confidence interval. A small but positive slope was found, consistent with stronger contextualization of words that are later in a sentence. Indices larger than 40 words are rare (0.8% of data) and are not plotted for visual clarity. B. ANOVA analysis of impact of Part of Speech on contextual representation. Black bars represent 95% confidence intervals. C. Similar to B but for semantic categories.

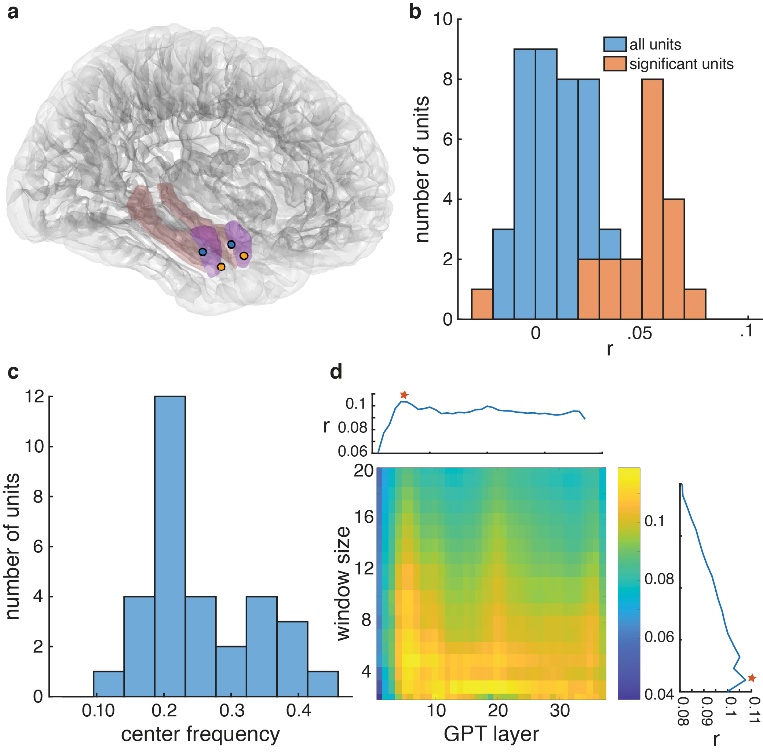

Extended Data Fig. 9: Position and Contextual Effects in the Amygdala A. Location of microelectrode recordings in two patients with bilateral amygdala recordings. Hippocampus is outlined in orange, amygdala in purple. B. Distribution of correlations between firing rate and word index in a sentence for all (blue) and significant (red) units. C. Distribution of center frequencies of units with significant sinusoidal encoding. D. Average match across all words between neural APG and GPT-2 APG for all layers (x axis) and window lengths from 2 to 20 (y axis). Average values projected above and to the right. Red stars reflect the maximums.

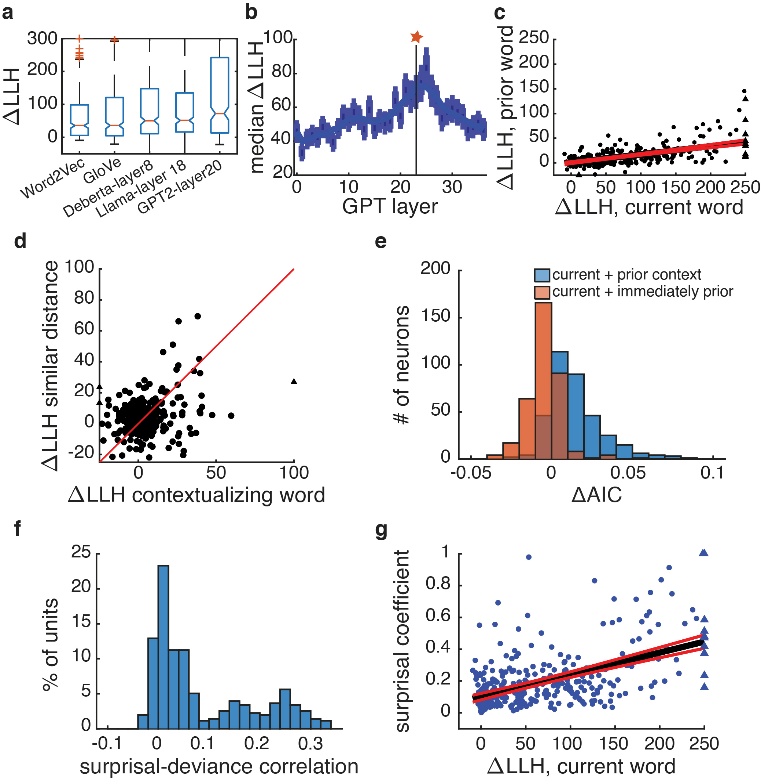

Extended Data Fig. 10: Reproduction of Results using Alternative Model Choices. A. A comparison of the LLH for each neuronal model with different semantic embeddings for non-contextual (Word2Vec and GloVe) versus contextual (DeBerta V3, Llama-3 and GPT-2). B. Box and whisker (median and SEM) plots of the distribution of LLH differences across all neurons using the hidden states of each layer of GPT-2 as the semantic predictors. Blue line represents the median smoothed with a gaussian of width 5 and star indicates maximum of fitted line. C. Plot of ΔLLH for current word versus next word for each neuron. Triangles are datapoints that are outside the limits of the plot. Black and red lines represent the best fit +\- 95% confidence interval. D. Plot of the model fits (as defined by the difference in LLH) for models that use contextualizing words versus models that use prior words with a similar distance distribution. The diagonal line represents the hypothesis that there is no difference, consistent with a general effect of autocorrelation. E. Histogram of the difference in the AIC per word between the basic current word model and the current + prior word model (blue) or the current + contextual word model (red). Positive values indicate that the unit’s firing patterns are better explained with both words, even when penalizing for the increased number of parameters. G. Distribution of correlations between the deviance of the model for each word and the relative surprise of that word for all neurons for all units. G. The ΔLLH of the non-contextual model as a function of the coefficients of surprisal in the linear model that predicts deviance. Note that subplots are positioned to match the locations in Figure 4.
